# Age-linked suppression of lipoxin A4 mediates cognitive deficits in mice and humans

**DOI:** 10.1101/2021.11.12.468379

**Authors:** Fabricio A. Pamplona, Gabriela Vitória, Felipe C. Ribeiro, Carolina A. Moraes, Pitia Flores Ledur, Karina Karmirian, Isis M. Ornelas, Luciana M. Leo, Bruna Paulsen, Felipe K. Sudo, Gabriel Coutinho, Claudia Drummond, Naima Assunção, Bart Vanderborght, Claudio Canetti, Hugo C. Castro-Faria-Neto, Paulo Mattos, Sergio T. Ferreira, Stevens K. Rehen, Fernando A. Bozza, Mychael V. Lourenco, Fernanda Tovar-Moll

## Abstract

Age increases the risk for cognitive impairment and is the single major risk factor for Alzheimer’s disease (AD), the most prevalent form of dementia in the elderly. The pathophysiological processes triggered by aging that render the brain vulnerable to dementia involve, at least in part, changes in inflammatory mediators. Here we show that lipoxin A4 (LXA4), a lipid mediator of inflammation resolution known to stimulate endocannabinoid signaling in the brain, is reduced in the aging central nervous system. We demonstrate that genetic suppression of 5-lipoxygenase (5-LOX), the enzyme mediating LXA4 synthesis, promotes learning impairment in mice. Conversely, administration of exogenous LXA4 attenuated cytokine production and memory loss induced by inflammation in mice. We further show that cerebrospinal fluid LXA4 is reduced in patients with dementia and positively associates with memory performance, brain-derived neurotrophic factor (BDNF), and AD-linked amyloid-β. Our findings suggest that reduced LXA4 levels may lead to vulnerability to age-related cognitive disorders and that promoting LXA4 signaling may comprise an effective strategy to prevent early cognitive decline in AD.

## Introduction

Inflammation constitutes an essential defensive mechanism that is dysregulated in aging and in several chronic and neurodegenerative disorders, including Alzheimer’s disease (AD) ^1, 2^. Accordingly, a chronic imbalance favoring disproportionate pro-inflammatory responses in the brain and periphery are thought to mediate age-related cognitive decline and AD pathogenesis ^3-9^.

Proper control of inflammatory responses depends on resolution processes, which are controlled by lipid mediators known as resolvins, such as lipoxins ^10^. Lipoxin A4 (LXA4) is an eicosanoid derived from arachidonic acid through transcellular metabolic pathways that depend on the enzymatic activity of 5-lipoxygenase (5-LOX) ^10-12^. In the periphery, LXA4 activates the G-protein-coupled receptor ALX to modulate gene expression towards resolution of inflammation and to promote immune cell recruitment to the site of infection or damage ^10^. Putative effects of LXA4 in the brain have long remained elusive, due to the low expression of ALX receptors in the central nervous system (CNS) ^13^.

We and others have suggested that LXA4 protects the central nervous system against injuries ^14-18^, including amyloid-β (Aβ) toxicity in mice ^15, 19, 20^. However, while the processes that propagate cytokine-mediated inflammation in AD have been thoroughly investigated, very little is known about the impact and mechanisms of LXA4 on brain function. Given that ALX receptors are poorly expressed in the central nervous system ^13^, other mechanisms should be responsible for the brain actions of LXA4. We have previously demonstrated that LXA4 binds to CB1 receptors in the brain to stimulate endocannabinoid signaling ^19^. These findings raise the prospect that LXA4 might be relevant to complex brain functions and that LXA4 signaling might be impaired in age-related cognitive diseases linked to aberrant inflammation, such as AD.

Herein we report that mammalian neurons and microglia produce considerable levels of LXA4. We further demonstrate that brain and peripheral LXA4, as well as memory, decline with aging and that genetic suppression of 5-LOX results in memory impairment in mice. Conversely, administration of exogenous LXA4 attenuated cytokine production and memory loss induced by an inflammatory stimulus in mice. We finally present results from a human study demonstrating that cerebrospinal fluid (CSF) LXA4 levels are reduced in dementia. Furthermore, LXA4 correlates with memory performance, Aβ and brain-derived neurotrophic factor (BDNF) in the human CSF. Altogether, these results support a protective role for LXA4 in the brain that is dysregulated in aging and, to a greater extent, in dementia.

## Methods

### Ethics

All mouse experiments were performed under protocols approved and supervised by the Institutional Animal Care and Use Committee of Federal University of Rio de Janeiro (CEUA-UFRJ) (protocol no. IBQM058). Experimental procedures involving human clinical data and CSF samples were approved by the Institutional Review Board (IRB) of Copa D’Or Hospital (protocol #47163715.0.0000.5249). Written informed consent was obtained from all volunteers. Samples were anonymized and measurements were performed by trained investigators in a blinded fashion. All studies have been performed according to the national and international ethical regulations and standards.

### iPSC cultures

Three different lines of human induced pluripotent stem cells were employed in this study - GM23279A iPSC line, obtained commercially from Coriell Institute Biobank; CF2 line, generated from fibroblasts; and C15 line, obtained from urine cells. Cells were reprogrammed using the CytoTune 2.0 Sendai Reprogramming kit (Thermo Fisher Scientific, USA) and characterization of the reprogrammed cell lines was conducted by immunostaining of both iPSCs colonies and iPSCs-derived embryoid bodies for self-renewal and three germ layer markers as described ^21, 22^. iPSCs were used to generate human neural progenitor cells (NPCs) by induction with retinoic acid (RA) for 18 days ^23, 24^. When neural tube-like structures emerged, they were collected and re-plated on adherent dishes treated with 10 µg/mL Poly-L-ornithine and 2.5 µg/mL laminin (Thermo Fisher Scientific). After passages using Accutase, homogenous cultures were obtained ^25^. NPCs were maintained and grown in DMEM/F-12 supplemented with N2, B-27, 25 ng/mL bFGF and 20 ng/mL EGF (Thermo Fisher Scientific), with medium changes every other day. The astrocyte cultures and mixed neuronal cultures were obtained from the differentiation of neural stem cells (NSCs), as described ^26^.

### Primary cell cultures

Primary astrocyte, microglial and neuronal cultures were obtained from Swiss mice. Brain tissue was dissociated into single cells in DMEM-F12 (Invitrogen; Carlsbad, CA, USA) supplemented with glutamine (2 mM), penicillin and streptomycin (`0.5 mg/mL, HyClone`), amphotericin B (0.65 μM, Sigma Aldrich, St. Louis, MO, USA) and 10% fetal bovine serum (FBS, Invitrogen; Invitrogen, Carlsbad, CA, USA). Cells were plated in 25 cm^2^ bottles pre-treated with poly-L-lysine (Sigma Aldrich, St. Louis, MO, USA). To generate astrocytes, cultures were maintained at 37 °C in an atmosphere of 5 % CO2 for 7 days until confluence was achieved. Confluent cells were passaged to generate purified astrocyte cultures. Microglia were isolated at day 13 *in vitro* by shaking for 45-60 minutes and plated on adherent dishes supplemented with DMEM-F12 with 10% FBS and maintained for more 24 hours. Hippocampal neurons were obtained from 16-day embryonic mice and maintained in Neurobasal medium (Invitrogen; Carlsbad, CA, USA) supplemented with B-27 (Life Technologies, Carlsbad CA, USA), penicillin, streptomycin (`0,5 mg/mL, HyClone`), glutamine (2×10-3 M), and fungizone (0.65 μM, Sigma Aldrich, St. Louis, MO, USA). Cultures were maintained at 37°C in a humidified atmosphere with 5% CO^2^ at 9 days in vitro. Cell homogenates were collected, centrifuged at 10000 g for 5 min at 4^°^C, and used for LXA4 measurements.

### Brain organoids

Brain organoids were generated from GM23279A induced pluripotent stem cell (iPSC) line as previously described ^27^. Briefly, 9,000 iPSC per well were plated of an ultra-low attachment 96-well plate in hESC medium (20% knockout serum replacement; Life Technologies) containing 50 µM ROCKi (Y27632; Merck Millipore, USA) and 4 ng/ml b-FGF. The plate was spun, and cells kept for 7 days to allow formation of embryoid bodies (EBs). On the seven following days, EBs were changed to ultra-low attachment plates, submitted to a Matrigel bath for 1h and media changed from neuroinduction media [1% N2 supplement (Gibco), 1% GlutaMAX (Life Technologies), 1% MEM-NEAAs, 1% P/S, and 1 μg/ml heparin in DMEM/F12 (Life Technologies)] to differentiation media minus vitamin A (50% neurobasal medium, 0.5% N2, 1% B27 supplement without vitamin A, 1:100 2-mercapto-ethanol, 0.5% MEM-NEAA, 1% GlutaMAX, and 1:100 P/S in DMEM/F12). On day 15, organoids were transferred to agitation in 6-well plates at 90 rpm, and media was changed to differentiation media plus vitamin A (50% neurobasal medium, 0.5% N2, 1% B27 supplement with vitamin A, 1:100 2-mercapto-ethanol, 0.5% MEM-NEAA, 1% GlutaMAX, and 1:100 P/S in DMEM/F12), and replaced every four days until ready for experiment. On day 45, brain organoids were fixed overnight in 4% PFA, then dehydrated in 30% sucrose, frozen in optimal cutting temperature compound on dry ice, and sectioned (20 μm thickness) with a cryostat (Leica Biosystems, Germany).

### Mouse strains

Swiss albino mice were provided by Fundação Oswaldo Cruz (FIOCRUZ); inbred C57BL/6 were provided by Charles River and 5-LOX knockouts (and wild-type littermates) were provided by FIOCRUZ and kept in the animal facilities at the Federal University of Rio de Janeiro. Adult male mice were used throughout the study and tested during the light phase of the light cycle. Mice were maintained on a 12 h light/dark cycle with food and water *ad libitum*, with maximum of 5 mice per cage.

### *In vivo* treatments

Adult male Swiss mice received an intraperitoneal (i.p.) injection of 0.3 mg/kg lipopolysaccharide (LPS) or vehicle immediately after contextual fear conditioning training session. One hour later, mice received an intracerebroventricular (i.c.v.) injection of either LXA4 (1 pmol) or vehicle and were tested for memory 7 days later.

### Immunofluorescence

For immunofluorescence in brain organoids, frozen sections were rinsed with PBS and permeabilized with PBS-Triton 0.3% (PBST) for 15 min. Slides were then incubated with 0.01M citrate buffer with 0.05% Tween 20 pH 6 for 10 min at 98°C for antigen retrieval and blocked with blocking solution (PBS with 3% BSA) for 2 hours at room temperature. Primary antibodies against 5-LOX (Cayman Chemical, 1:100) or MAP2 (Sigma-Aldrich, 1:300) diluted in blocking solution were incubated at 4°C overnight. Sections were then rinsed with PBS and incubated in AlexaFluor-conjugated IgG secondary antibody goat anti-mouse, rabbit or goat (Invitrogen, 1:400) for 1 hour at room temperature and washed 3 times for 5 min in PBS. For nuclear staining, sections were incubated with DAPI for 5 min. Slides were washed again 3 times for 5 min in PBS and then cover-slipped with Aqua-Poly/Mount (Polysciences, Inc). Images were acquired using a Leica TCS SP8 confocal microscope. For immunofluorescence experiments in the mouse hippocampus, mice were transcardially perfused with saline and 4% formaldehyde. Brains were removed and post-fixed in 4% formaldehyde overnight, then submerged in 30% sucrose solution. Forty μm-thick hippocampal cryosections were obtained, mounted on glass slides, and exposed to 0.01M citrate buffer for antigen retrieval for 20 min at 60°C. Tissue was permeabilized with 0.5% Triton X-100 in 50 mM ammonium chloride for 20 min at room temperature. Sections were then incubated in blocking solution (5% bovine serum albumin in PBST) for 60 min at room temperature and then incubated with anti-CB1R (Merck Millipore, 1:1000) in blocking solution overnight at 4 °C. Sections were then rinsed with PBST and incubated with AlexaFluor-conjugated anti-rabbit IgG secondary antibodies (1:400) for 2 hours at room temperature, followed by a short 5 min incubation in a DAPI solution (Molecular Probes). A 5 min incubation in 1% Sudan Black B prepared in 70% ethanol was applied to quench autofluorescence. Tissue was mounted on coverslips with Aqua Poly/Mount (Polysciences) and imaged on a Zeiss AxioImager M2 microscope. Image fluorescence was quantified on ImageJ ^28^ after selection of the stratum pyramidale and stratum radiatum of hippocampal CA1 and CA3 subfields as regions of interest.

### Human samples

CSF samples for this study were obtained from a cohort recruited at the D’Or Institute of Research and Education (IDOR) in Rio de Janeiro, Brazil. This cohort comprised healthy controls (HC; N = 25), and cognitively impaired subjects with diagnosis of either amnestic mild cognitive impairment (aMCI; N = 13), Alzheimer’s disease (N = 14), or dementia with Lewy bodies (DLB; N = 9). Inclusion criteria for this study were: age > 60 years; absence of other neurological conditions, neurodevelopmental or genetic diseases; native Brazilian Portuguese speakers; formal education ≥ 8 years; no restriction for MRI studies; no severe metabolic disease). All patients were evaluated with the same extensive clinical, neuropsychological and neuroimaging investigation as described ^29-33^. For demographics and biomarker information, see Table□1. CSF samples were collected by lumbar puncture performed around 11 a.m. in all cases, to minimize circadian fluctuations. CSF was centrifuged, and the supernatant was collected, aliquoted, and immediately frozen at −80°C. Before assays, samples were thawed and kept on ice until use. Samples and calibrators were run in duplicates.

### ELISA

Mouse brains were harvested, and lipid extraction was perfomed with ethanol (5 μL/mg of wet tissue) followed by centrifugation for 5 min at 10,000 × g. The supernatant was applied directly into a double-sandwich ELISA kit, read at 650 nm and normalized by wet tissue weight (g). Plasma and CSF samples and culture supernatants were applied to the ELISA kits according to manufacturer’s instructions. ELISA kits for LXA4 (#EA46) were from Oxford Biomedical Research (Rochester Hills, MI). ELISA kits for Aβ_42_, and total tau (t-tau) were provided by Euroimmun (Lu□beck, Germany) and experiments were run following manufacturer’s instructions.

### Inhibitory avoidance

The step-down inhibitory avoidance apparatus consisted of a box measuring 26 × 10 × 35 cm with a 10 × 10 × 4 cm platform placed in the center, surrounded by a floor made of parallel bronze bars and connected to a power source. For the training sessions, mice were placed on the platform and when they stepped down with four paws onto the grid they received a 0.7 mA foot shock for 2 s, and immediately returned to their home cage. For the test session (30 min or 24 h after training), animals were again placed on top of the platform, and latency to step down was recorded. For the protocol with multiple trials, a milder shock of 0.5 mA was used, and the procedure was repeated until the animal learned to remain 180 s on the platform for 180 s (criteria). For memory extinction, mice were conditioned in the step-down inhibitory avoidance task, as described above, only with a 1 mA foot shock intensity. Starting on the 12^th^ day after conditioning, mice were submitted to successive extinction trials in 24 h intervals, where they were placed on the platform and allowed to step down onto the grid in the absence of a foot shock. The latency to step down was recorded and mice were allowed to freely explore the apparatus for 30 s after every trial. Reduction in step-down latency across successive trials indicated extinction learning.

### Contextual Fear Conditioning

To assess contextual fear memory consolidation, a two-phase protocol was used. Swiss mice were initially trained in the conditioning cage (40 × 25 × 30 cm) and were allowed to freely explore for 3 minutes followed by application of a single foot shock (1 mA) for 2 s. Mice were kept for another 1 minute in the cage and removed. Right after training, mice received an intraperitoneal injection of 0.3 mg/kg lipopolysaccharide (LPS) obtained from E. coli 0127:B8 (Sigma-Aldrich) or vehicle. One hour later, mice received an intracerebroventricular (i.c.v.) injection of either LXA4 (1 pmol) or vehicle. Seven days after training, mice were presented to the same cage for 5 minutes without receiving a foot shock. Freezing behavior was recorded automatically using the Freezing software (Panlab; Cornella, Spain). In all behavioral experiments, the experimenter was blinded to the groups tested.

### Statistical analysis

Statistical analyses were performed using GraphPad Prism 6 software (La Jolla, CA) or the IBM SPSS Statistics v. 26 (Armonk, NY). Differences between two independent groups were analyzed using Student’s t-test. When three or more independent experimental groups were compared, one-or two-way ANOVA was used, followed by appropriate *post hoc* tests, as stated in “Figure Legends”. For correlations, data distribution was initially assessed using Shapiro-Wilk Test. After adjustment for age, partial rank correlations were performed to assess the relationship between CSF LXA4 levels and clinical/biomarker variables. Significance level was set at 0.05. The predictive power of CSF LXA4, alone or in combination with Aβ_42_, to identify clinical AD was tested by plotting receiver operating characteristic (ROC) curves, widely used to determine the diagnostic potential of biomarkers, and determining the area under the curve (AUC) with a confidence interval of 0.95.

## Results

### LXA4 is produced by neurons and microglia

Lipoxins, including LXA4, are locally produced by immune cells at sites of inflammation or systemically ^11^. Although we and others have previously reported that LXA4 is bioactive in the brain ^14, 15, 19^, whether brain cells comprise a source for LXA4 remains unknown. We first used primary cultures to investigate whether brain cells produced LXA4 at detectable levels. We found that mouse hippocampal neurons and microglia had significantly higher content of LXA4 than astrocytes (Fig. 1a). We next used human induced pluripotent stem cells (iPSCs) to derive human neural progenitor cell (NPC), neuron, astrocyte or microglial-like cultures to determine the levels of LXA4. We found that human iPSC-derived neurons produce considerably higher levels of LXA4 than astrocytes or NPCs (Fig. 1a). Furthermore, in line with mouse results, levels of human microglia-sourced LXA4 are comparable to human neurons, suggesting microglia as a key producer of LXA4 (Fig. 1a). To further ascertain whether neuronal 5-LOX expression would be present in more complex systems, we performed immunohistochemical experiments to label 5-LOX in 45-day-old human brain organoids derived from iPSC. At that stage, human brain organoids express a biochemical profile, patterning, and structural organization similar to what is observed in the human embryonic brain ^27^. We observed prominent 5-LOX labeling in microtubule-associated protein 2 (MAP2)-positive cells (Fig. 1b), confirming that human neurons typically located in the outer layers of human brain organoids express 5-LOX. Together, our results raise the possibility that neurons and microglia are important sources of LXA4 in the human brain.

**Figure 1.**
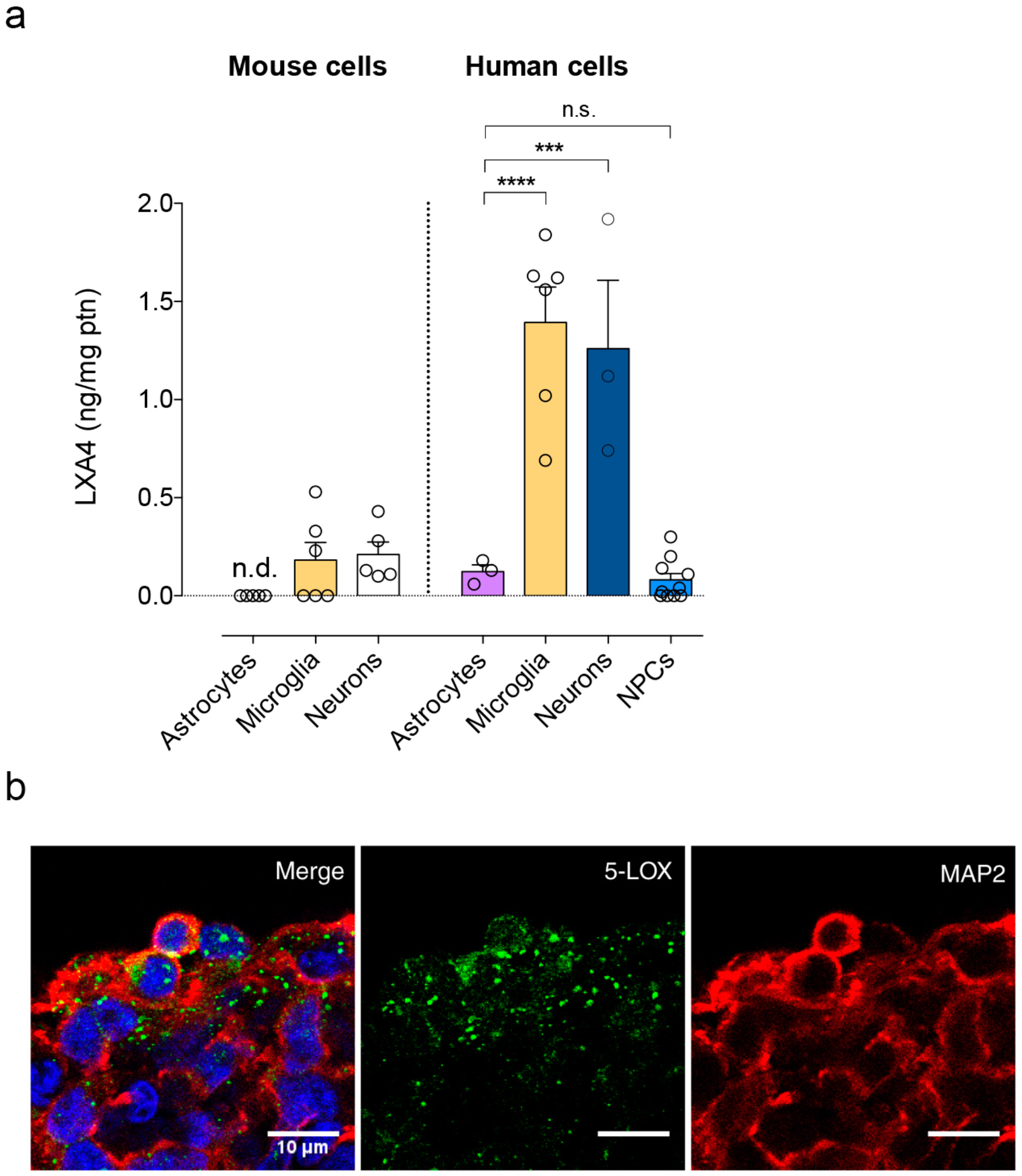
LXA4 is produced by neurons and microglia. (a) Levels of lipoxin A4 (LXA4) in primary cultures enriched in astrocytes, microglia or neurons derived from mice (left) or in human iPSC-derived astrocytes, microglia, neurons, or neural progenitor cells (NPCs). Unpaired two-tailed one-way ANOVA with Šidák post hoc test (*** p < 0.001; **** p < 0.0001; n.s. non-significant). n.d.; not detected. (b) Representative images of immunofluorescence experiments (5-lipoxygenase, 5-LOX immunoreactivity: green; microtubule-associated protein 2, MAP2 immunoreactivity: red; DAPI: blue; N=3) in human iPSC-derived brain organoids. Scale bar: 10 μm.

### Aging reduces systemic and brain LXA4 levels in mice

Since aging impairs systemic response to injuries ^34, 35^ and renders the brain vulnerable to neurodegenerative conditions ^36^, we next determined whether levels of LXA4 would be modified by aging in mice. We found that plasma and brain LXA4 levels were reduced in 12-month-old male Swiss mice compared to young 3-month-old controls (Fig. 2a,b). This is accompanied by evident short-term memory impairment in the inhibitory avoidance memory test, as aged mice presented reduced latency to step down from the platform in the test sessions (Fig. 2c). Results indicate that reduced brain and peripheral LXA4 levels concur with cognitive deficits in aged mice.

**Figure 2.**
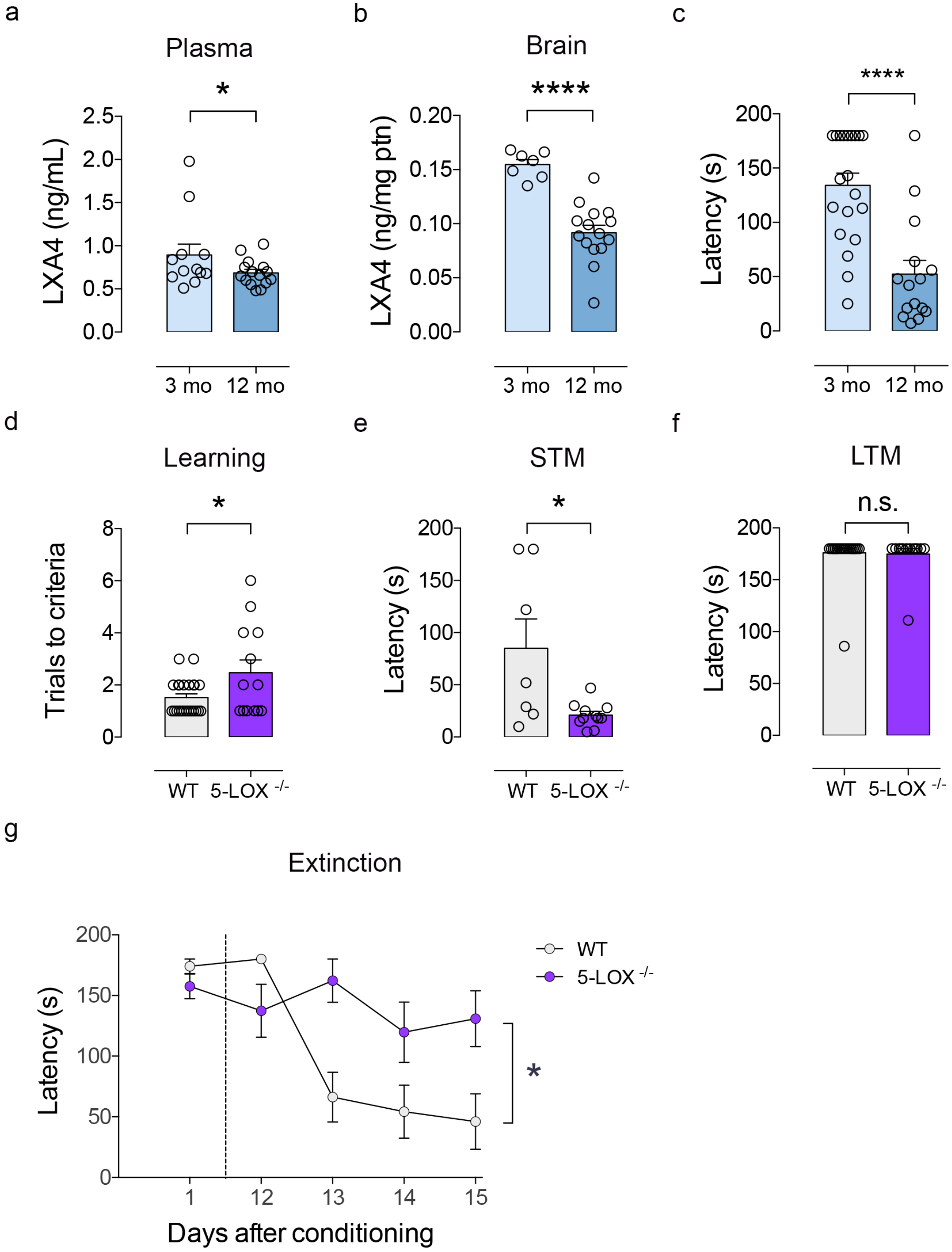
Age-linked reductions in LXA4 result in cognitive impairment. (a,b) Levels of lipoxin A4 (LXA4) in plasma (a) and brain (b) of young (3 month-old) or aged (12 month-old) Swiss mice. (c) Step-down latency of young or aged mice in the inhibitory avoidance task. (d) Number of trials required for each mouse to reach the criteria during the training session in the inhibitory avoidance fear task. (e,f) Step-down latency of 5-LOX^-/-^ or WT mice (3 month-old) in the inhibitory avoidance task to assess short-term (e; 1 hour after training) or long-term memory (f; 24 hours after training). For *a, b*, and *d*, two-tailed unpaired Student’s t-test. For *c, e*, and *f*, two-tailed unpaired Mann-Whitney (* p < 0.05; **** p < 0.0001). Graphs show means ± standard error of the mean (SEM). Each dot represents an individual. (g) Fear memory extinction assessed by step-down latency of 5-LOX^-/-^ or WT after repeated test sessions twelve to fifteen days after original conditioning session. Repeated measures one-way ANOVA with Šidák post hoc test (* p < 0.05). Graphs show means ± SEM.

### Suppression of LXA4 mimics age-associated memory loss in mice

We next hypothesized that reductions in LXA4 would be causally implicated in memory impairments in mice. Therefore, we tested memory performance in 3-month-old (young) 5-LOX homozygous knockout mice (5-LOX^-/-^), which show reduced circulating LXA4 levels ^19^, and their respective wild-type littermates (WT) in the inhibitory avoidance memory task. We used adult 5-LOX^-/-^mice in these experiments to mimic reductions in circulating LXA4 and specifically dissect the roles of 5-LOX, while avoiding potential confounders triggered by normal aging. Control experiments revealed no differences in body weight (Supp. Fig. S1a) or food and water intake (Supp. Fig. 1b,c) across genotypes. Additionally, levels of hippocampal CB1 receptors were similar in 5-LOX^-/-^ and WT mice (Supp. Fig. 1d-f).

5-LOX^-/-^ mice took significantly more learning sessions in the inhibitory avoidance task to reach criteria than WT (Fig. 2d). While WT showed normal memory performance, 5-LOX^-/-^ mice presented impaired short-term, but not long-term memory (Fig. 2e,f). Furthermore, 5-LOX^-/-^ mice had impaired extinction of learned fear memory (Fig. 2g), which is in further agreement with learning deficits. Altogether, these results demonstrate that 5-LOX-mediated LXA4 synthesis supports proper cognitive function and indicate that reductions in LXA4 may render the brain vulnerable to dysfunction and memory impairment.

### Administration of LXA4 attenuates inflammatory responses and memory loss

Our results so far indicated that LXA4-mediated signaling could be part of a protective mechanism against age-related cognitive decline, which is, at least in part, mediated by inflammation ^37^. We then sought to determine whether administration of LXA4 would protect against impairments in memory consolidation induced by an inflammatory injury. Mice were trained in a contextual fear conditioning paradigm and received a post-training intraperitoneal (i.p.) injection of 0.3 mg/kg lipopolysaccharide (LPS) or vehicle. One hour after LPS, mice received an intracerebroventricular (i.c.v.) injection of either LXA4 (1 pmol) or vehicle and were tested for memory 7 days later. We found that i.c.v. LXA4 administration rescued impaired contextual fear memory consolidation in LPS-injected mice (Fig. 3a), advocating towards a neuroprotective effect via reduction of inflammatory pathways.

**Figure 3.**
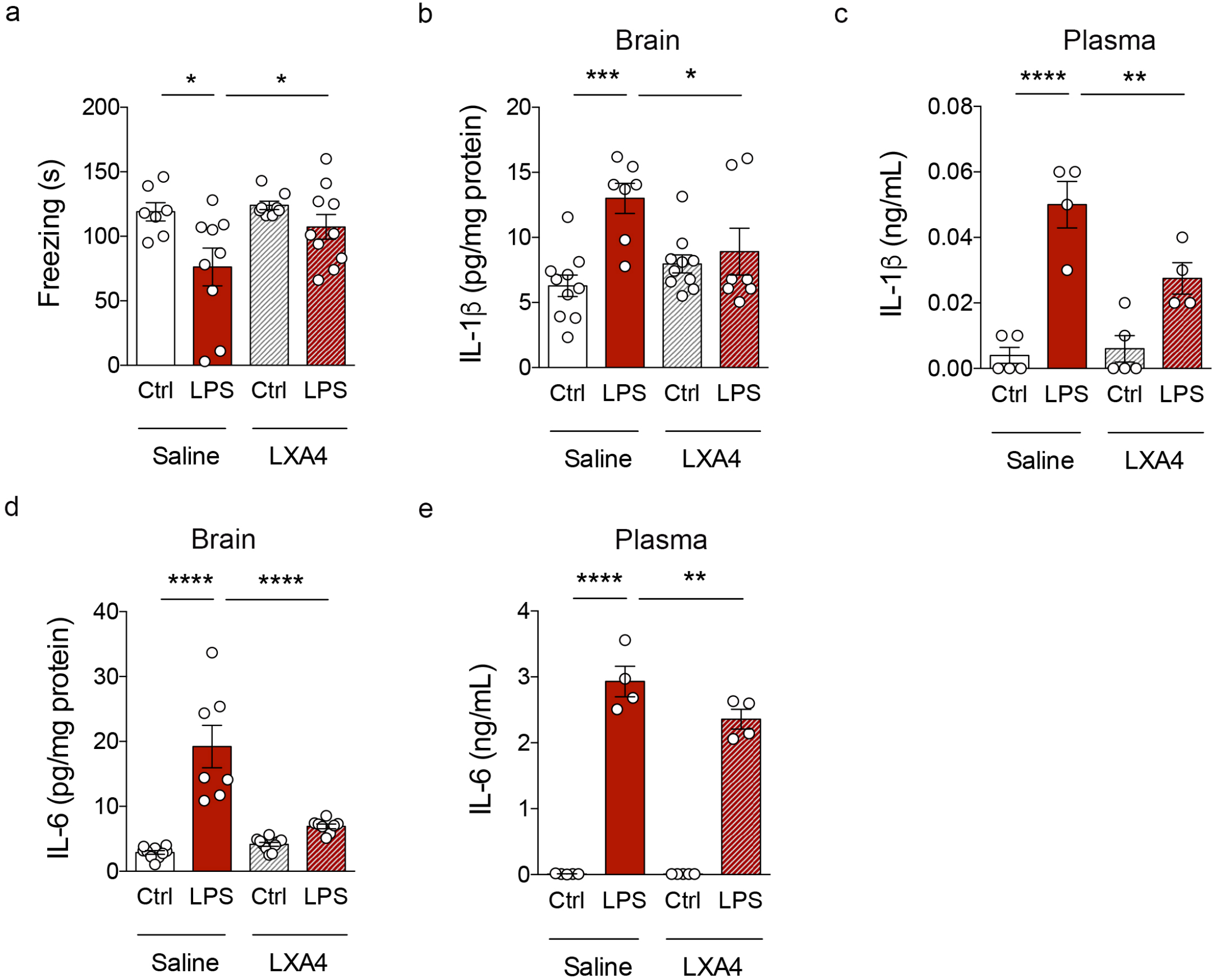
LXA4 attenuates inflammation-induced memory failure and cytokine production in mice. (a) Freezing (seconds) of control or LPS-injected mice (0.3 mg/kg) treated with vehicle or 1 pmol LXA4 (i.c.v) in the contextual fear conditioning task. (b-e) Brain and plasma levels of IL-1β (b,c) or IL-6 (d,e) in control or LPS-injected mice (0.3 mg/kg; i.p.) treated with vehicle or 1 pmol LXA4 (i.c.v). Two-tailed unpaired two-way ANOVA followed by Holm-Šidák post hoc test (* p < 0.05; ** p < 0.01; *** p < 0.001; **** p < 0.0001). Graphs show means ± standard error of the mean (SEM).

We thus attempted to determine the impact of LXA4 on LPS-induced inflammatory cytokine production. I.c.v. LXA4 administration reduced levels of interleukin 1β (IL-1β) (Fig. 3b,c) and interleukin 6 (IL-6) (Fig. 3d,e) in the brain and plasma of LPS-treated mice. These results indicate that LXA4 attenuates peripheral and brain inflammation, and rescues inflammation-induced cognitive defects in mice.

### Cerebrospinal fluid LXA4 declines with aging and dementia in humans

To test the potential relevance of LXA4 in humans, we investigated LXA4 levels in the cerebrospinal fluid (CSF) of a cohort of human subjects in a memory clinic. They were diagnosed as mild cognitive impairment (MCI), AD, dementia with Lewy bodies (DLB), or controls (see Supp. Table 1 for demographics and biomarker information). We initially found that LXA4 declines with aging in human CSF in this cohort (Fig. 4a), in line with our mouse studies. We further determined that LXA4 levels were markedly decreased in subjects with AD or DLB, but not in MCI, as compared to controls (Fig. 4b). Together, these results support the notion that aging leads to a reduction in brain LXA4 levels. The decrease in LXA4-mediated protective mechanisms may lead to vulnerability to cognitive diseases and, presumably, neurodegeneration.

**Figure 4.**
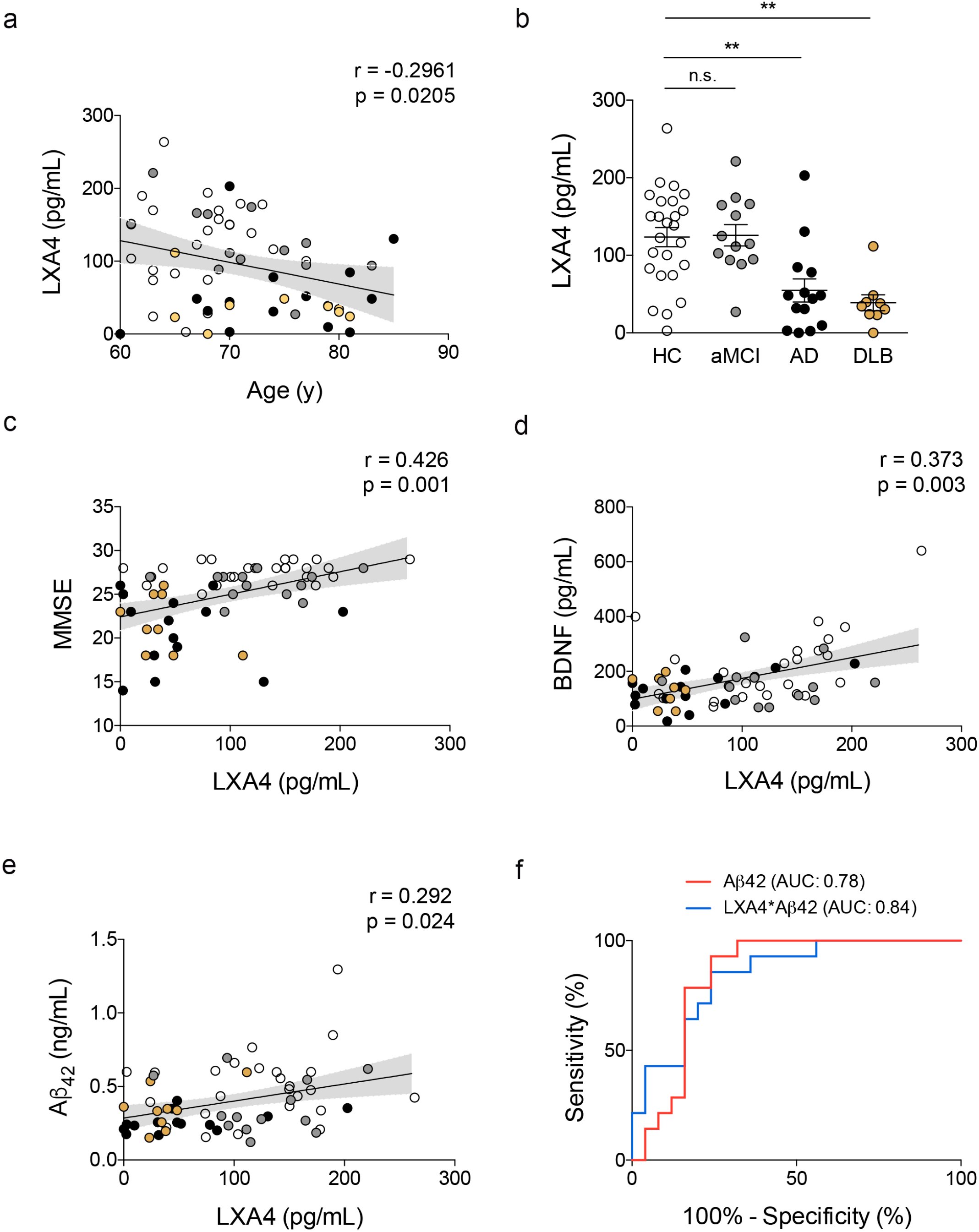
Cerebrospinal fluid LXA4 is reduced in aging and in dementia in humans. (a) Correlation between age (in years) and CSF LXA4 in human subjects. (b) CSF levels of LXA4 in AD and DLB patients compared to healthy controls or amnestic MCI patients (N = 25 controls, 13 aMCI, 14 AD, 9 LBD patients). Two-tailed unpaired one-way ANOVA followed by Holm– Šidák post hoc test (** p < 0.01; ns, non-significant). Graphs show means ± standard error of the mean (SEM). (c-e) Correlations between LXA4 and MMSE scores (c), CSF BDNF (d) or Aβ_42_ (e) levels in human subjects. Lines represent partial rank correlations (r and p-values as indicated in graphs), adjusted for age, and the confidence interval is represented as gray shade. (f) Receiver-operating characteristic curves for diagnostic based on Aβ_42_ alone (red line) or LXA4*Aβ_42_ (blue line); confidence interval: 0.95; p < 0.001. HC: healthy controls, white symbols; aMCI: amnestic mild cognitive impairment, grey symbols; AD: Alzheimer’s disease, black symbols; DLB: dementia with Lewy bodies, golden symbols); AUC: area under curve.

### Cerebrospinal fluid LXA4 correlates with memory performance, BDNF, and Aβ_42_ levels in humans

We then investigated whether CSF LXA4 would associate with memory performance and the levels of markers relevant to AD. CSF LXA4 showed positive correlations with mini-mental state exam (MMSE) scores, a proxy for memory performance in humans (Fig. 4c). We further found that LXA4 positively correlates with BDNF, a neurotrophin essential for memory function ^38, 39^, in the CSF (Fig. 4d). These results indicate that central LXA4 actions may favor brain homeostasis and cognition.

We next addressed potential associations between CSF LXA4 and AD biomarkers (Aβ_42_ and tau). We found that CSF LXA4 positively associates with CSF Aβ_42_ (Fig. 4e), but not with tau (Supp.Fig. 2). Finally, combination of CSF LXA4 and Aβ_42_ slightly increased the sensitivity and specificity of AD diagnostics in a receiver operating characteristic (ROC) curve (Fig. 4f). These results suggest that declines in CSF LXA4 correlate with brain Aβ_42_ accumulation, possibly facilitating its neurotoxic actions.

## Discussion

Age-related cognitive impairment and dementia are amongst the causes of significant disability in the elderly and effective interventions are still not available. Nonetheless, it is well established that loss of immune homeostasis and aberrant inflammatory responses comprise significant factors predisposing individuals to brain dysfunction and dementia ^40, 41^. Here we provide evidence suggesting that LXA4 is part of an endogenous protective mechanism that decreases with aging and renders the brain vulnerable to memory failure and dementia.

We first addressed a long-standing question of whether brain cells produce LXA4. Previous evidence indicated that rodent neurons and microglia express 5-LOX, the enzyme required for LXA4 synthesis ^16, 17^. Nonetheless, direct evidence for LXA4 production in brain cells was lacking. We found that mouse and human neurons and microglia in culture actively produce and release LXA4, whereas astrocytes produce smaller amounts of LXA4. These results suggest that neurons and microglia could act as local sources of LXA4 in the brain and builds upon our previous data showing that LXA4 is present in the brain despite the absence of its canonical ALX receptor, at least under physiological conditions ^19^.

We and others have provided initial evidence suggesting that peripheral LXA4 levels are reduced with aging in mice and humans. While we have previously shown that 12-month-old mice present reduced plasma levels of LXA4 as compared to 3-month-old mice ^14^, Gangemi et al. (2005) demonstrated that urinary LXA4 is markedly reduced in aged humans ^34^. We now confirmed these previous results in plasma and extended our studies to demonstrate that central LXA4 also declines with age and accompanies memory impairment in mice.

To further support that reduced LXA4 underlies memory impairment, we determined that 5-LOX^-/-^ mice, which present reduced plasma and brain LXA4 content, present learning impairment in an inhibitory avoidance task. We decided to use 3-month-old 5-LOX^-/-^ mice (instead of aged subjects) in these experiments to circumvent a potential source of confusion with other aspects related to the aging process. For instance, 5-LOX^-/-^ mice have been reported to develop additional age-dependent behavioral alterations, such as anxiety-like behavior ^42^, which are not present in young mice ^14^. Together, these results demonstrate that reduced LXA4 is sufficient to trigger memory failure in mice. Future studies are warranted to determine the relative contributions of circulating and brain LXA4 to cognition by ablating 5-LOX in specific sources of LXA4 (i.e. neurons, microglia, neutrophil, platelets).

5-LOX was previously shown to have a deleterious impact in mouse models of AD and tauopathy ^43-46^, potentially due to dyshomeostasis of other lipid derivatives. Conversely, stimulation of LXA4 signaling through aspirin-triggered lipoxin (ATL) resulted in beneficial actions, including reduced microglial reactivity, less Aβ_42_ accumulation and tau phosphorylation, and attenuation of memory impairment in mouse models of AD ^15, 20^. Notably, LXA4 was reported to attenuate Aβ-induced memory impairment through CB1 receptors ^19^. Our findings further demonstrate that brain administration of LXA4 prevents inflammation-induced cytokine upregulation and memory consolidation defects in mice. Results suggest that preserving LXA4 signaling across adulthood might be beneficial to ward off persistent inflammation, cognitive decline, and AD risk at later stages of life.

We have previously shown that LXA4 acts as an allosteric agonist of CB1 receptors in the brain, enhancing the affinity of CB1 receptors to anandamide ^19^. Given that endocannabinoid signaling through CB1 receptors is essential for proper brain functions ^47-50^, locally produced LXA4 could serve as a means of intercellular communication in the CNS to fuel synaptic plasticity, potentiate astrocyte metabolism and microglial surveillance. Conversely, reduced brain LXA4 could explain, at least in part, compromised CB1 receptor signaling, which may then translate into memory impairment ^51^.

Our results show that CSF LXA4 is considerably reduced in patients with dementia (AD and DLB), conferring translational relevance to the rodent studies. Significant correlations of CSF LXA4 with BDNF and memory function in humans also indicate potential pro-mnemonic actions of central LXA4 and highlight the need for further translational investigation of the underlying mechanisms. An intriguing observation relies on the positive correlation of CSF LXA4 with Aβ_42_, but not with tau, in humans. This suggests that brain Aβ accumulation (as assessed by lower CSF Aβ_42_) may moderate LXA4 levels in the CNS independently of tau pathology (as determined by CSF tau). These results encourage additional investigation to clarify the interrelation of Aβ_42_, tau and LXA4, as well as to determine whether incorporation of LXA4 measurements into a CSF biomarker panel can aid in the discrimination of a subset of AD cases, thereby improving overall diagnostics.

Activation of the innate immune system appears to mediate cognitive decline and AD pathogenesis ^37, 52^. Indeed, aberrant cytokine release ^53, 54^ and abnormal innate immunity response to brain Aβ deposition have been reported in mouse models and patients of AD ^55, 56^. Infiltration of immune cells, such as neutrophils ^57^, into the brain may shift cellular responses towards reduced LXA4 production and, consequently, increased neurotoxicity in AD.

We hypothesize that LXA4 deploys a dual mechanism as an anti-inflammatory and neuroprotective agent, thereby contributing to resilience of central nervous system against disease-associated insults, such as Aβ_42_ in AD. Hence, LXA4 levels may be useful as a biomarker of brain vulnerability, and its decreased levels may indicate a homeostatic breakdown. Since a significant part of the CB1 receptor-related effects of anandamide are due to synergistic LXA4 interaction ^19^, the vulnerability window caused by age-associated LXA4 reduction might be conceived as part of the “endocannabinoid deficiency syndrome” hypothesis ^58^. This concept advocates that under circumstances of relatively low endocannabinoid signaling, certain inflammation-linked injuries, such as fibromyalgia, inflammatory bowel syndrome and migraine, become more incident. Here we propose to extend this concept to include neurodegeneration and dementia.

In summary, our findings support the notion that LXA4 signaling may comprise an endogenous protective mechanism that renders the brain vulnerable to the exacerbated inflammation and neurodegenerative factors (i.e. Aβ_42_ deposition). Preventing declines in LXA4 levels may preserve endocannabinoid signaling, brain homeostasis, and cognition, thereby contributing to reduced odds of dementia. Future studies are needed to establish the intricate signaling pathways initiated by brain actions of LXA4, as well as to test whether stimulation of LXA4 (or the endocannabinoid system) is effective in delaying cognitive decline in aged humans.

## Supporting information

Suppplemental Material

## Acknowledgements

This work was supported by grants from Fundação Carlos Chagas Filho de Amparo à Pesquisa do Estado do Rio de Janeiro (FAPERJ) to FAP, HCCFN, and MVL, Conselho Nacional de Desenvolvimento Científico e Tecnológico (CNPq) to FAP and MVL, from Alzheimer’s Association (AARG-D-615714) to MVL, Serrapilheira Institute (R-2012-37967) to MVL, and intramural grants from the D’Or Institute for Research and Education (IDOR) and Rede D’Or São Luiz Hospital Network. We thank the IDOR clinical and imaging staff for patients’ recruitment and diagnostic investigation.

## Author contributions

FAP, FTM designed the study. FAP, MVL, FCR, GV, CM, PFL, KK, IMO, LML, BP, FKS, GC, CD, NA, BV, and CC performed research. FAP, MVL, BP, and FKS analyzed data. FAP, MVL, CC, HCCFN, SKR, FAB, and FTM contributed reagents, materials, animals, and analysis tools. FAP, MVL, STF, FAB, and FTM analyzed and discussed results. FAP and MVL wrote the manuscript.

