## Supplementary material for "Age-linked suppression of lipoxin A4 mediates cognitive deficits in mice and humans": Suppplemental Material

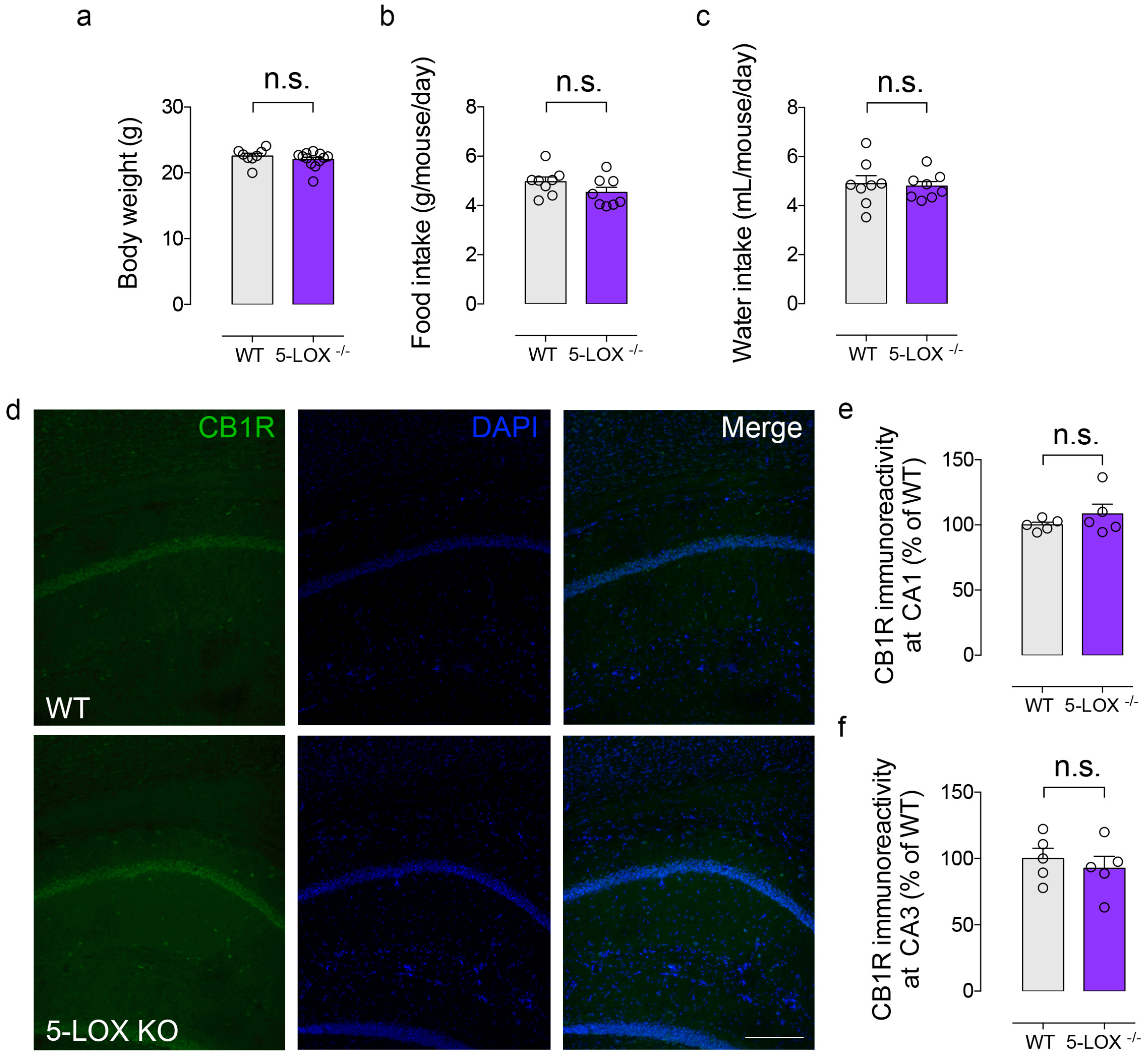


**Supplemental Figure 1. 5-LOX$^{-/-}$ mice do not present alterations in body weight, food/water intake, or hippocampal endocannabionoid receptor 1 (CB1R) expression.** (a-c) Body weight (a), 24-hour food intake (b) and 24-hour water intake (c) in adult male 5-LOX$^{-/-}$ or WT mice. (d) Representative images of immunofluorescence experiments (CB1R immunoreactivity: green; DAPI immunoreactivity: blue) in the hippocampal formation of 5-LOX$^{-/-}$ or WT mice. Scale bar: 50 µm. (e,f) Quantification of CB1R immunoreactivity at CA1 (e) and CA3 (f) hippocampal subregions obtained from experiments in *d* (n = 5 mice per group) Unpaired two-tailed Student's t-test; ns, non-significant. Graphs show mean ± standard error of the mean (SEM).

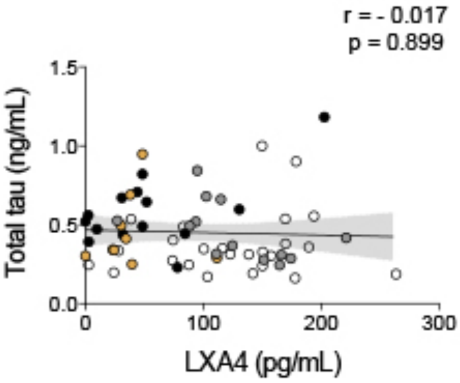


**Supplemental Figure 2. LXA4 does not correlate with total tau levels in the CSF.** Line represents a partial rank correlation (r and p-values as indicated in graph) and the confidence interval is represented as gray shade.

**Supplementary Table 1. Demographic, clinical and biomarker characteristics of donor subjects**

| | Healthy controls (HC) | Mild cognitive impairment (MCI) | Alzheimer's disease (AD) | Dementia with Lewy bodies (DLB) | F, $\chi^2$ (p-value) |
|---|---|---|---|---|---|
| Sex, male/female | 10/15 | 8/5 | 4/10 | 2/7 | 4.5 (0.21) |
| Age (years) | 67.8 ± 4.8 (61-79) | 71.5 ± 6.1 (61-83) | 74.2 ± 7.1* (60-85) | 73.7 ± 6.7 (65-81) | 4.4 (0.01) |
| Antidepressant use, yes/no | 6/19 | 6/7 | 6/8 | 6/3 | 5.6 (0.13) |
| BMI | 26.6 ± 5.0 (20-40.8) | 27.1 ± 3.1 (23.7-33.1) | 26.3 ± 4.4 (20.6-35.8) | 27.7 ± 6.0 (20-40.3) | 0.2 (0.90) |
| MMSE | 27.6 ± 1.2 (25-29) | 26.1 ± 1.5 (23-28) | 20.9 ± 4.1**** (14-26) | 21.7 ± 3.2**** (18-26) | 39.9 (<0.0001) |
| ApoE4, positive/ negative | 7/18 | 6/7 | 6/8 | 5/4 | 2.7 (0.44) |
| CSF A$\beta_{42}$ (pg/mL) | 507 ± 242 (156-1296) | 364 ± 185 (122-695) | 260 ± 68.7** (169-403) | 347 ± 145 (152-597) | 12.44 (0.002) |
| CSF t-tau (pg/mL) | 374 ± 207 (162-999) | 457 ± 184 (244-844) | 583 ± 227*** (230-1,181) | 452 ± 229 (250-947) | 10.5 (0.01) |

Values are presented as means ± SD (range). Statistical significances presented as F (p-value) based on two-tailed one-way ANOVA followed by Holm-Sidak adjustment for multiple comparisons, except for sex and APOE4 (Chi-Square Test, $\chi^2$ (p-value); A$\beta_{42}$ and tau (Kruskal-Wallis test followed by Dunn's adjustment for multiple comparisons). Asterisks indicate statistically significant differences from HC (* $p < 0.05$; ** $p < 0.01$; *** $p<0.001$; **** $p < 0.0001$).

Abbreviations: $A\beta_{42}$, amyloid-$\beta_{1-42}$; AD, Alzheimer's disease; APOE4, apolipoprotein E4; CSF, cerebrospinal fluid; DLB, Dementia with Lewy bodies; HC, Healthy controls; MCI, Mild cognitive impairment; MMSE, Mini-Mental State Exam; t-tau, total tau.
